# Improvement of beef cattle cow’s pregnancy rate using an effective dose of Galectin-1

**DOI:** 10.1101/2021.12.07.471574

**Authors:** Erika da Silva Carvalho Morani, Helen Alves Penha, Fernando Sebastián Baldi Rey, Marcelo Rácoletta

## Abstract

Galectins are mentioned in the literature as multifunctional molecules that participate in several biological processes such as adhesion, cell proliferation, and cycle, apoptosis, RNA processing, inflammatory process control, and reproductive physiological mechanisms. Galectin-1 has been referred to as a mediator involved in the prevention of early embryonic mortality in mammals. Exogenous GAL-1 (eGAL-1) can be found in Tolerana®. The objective of the study was to evaluate whether eGAL-1 can increase the pregnancy rate when used in an AI procedure (in a complementary artificial insemination procedure, using a second AI gun). The pregnancy rate was determined by the pregnancy condition through an ultrasound exam performed 25 to 35 days after the fixed-time artificial insemination (FTAI) of breeding cows (n=3,469 beef cows). The efficacy of GAL-1 was evaluated by comparing the pregnancy rate of the two groups (Treatment and Control Groups) in 107 contemporary groups (YG) established by the created statistical model. Based on the obtained results, it can be confirmed that the correct administration of a single dose of eGAL-1 can increase the probability of obtaining pregnancy in beef cows by up to 8.68% (p < 0.0001). The recommendation of the use of eGAL-1 during the FTAI procedure was reasonable in the beef cattle AI routine. On average, the complete procedure, using eGAL-1, took about 10 ± 5 seconds more time than the conventional procedure.

## Introduction

Cattle pregnancy loss can be considered one of the major challenges in the handling and cause of economic losses to the producer.

The average pregnancy rate in beef cattle, considering a single service, varies widely in the available literature. Now the fertilization rate in beef cattle can be high, reaching up to 95% in some scenarios, however, calculating the pregnancy rate by considering the complete breeding season, where 2 to 3 Artificial Insemination (AI) procedures and more exposure to the bulls are carried out. Embryonic mortality occurs mainly during the first 30 days of pregnancy, considerably reducing reproductive efficiency (Diskin and Morris, 2008; Diskin et al., 2016). Estimates indicate that pregnancy loss can reach 48% with one insemination during this period (Reese et al., 2020), which compromises the possibility of obtaining high pregnancy rates.

Several causes explain the reduction in the pregnancy rate, including early embryonic mortality (Diskin and Morris, 2008); diseases (Cheng et al., 2016), uterine asynchrony inducing failure of maternal recognition of pregnancy (Pope, 1988; Pohler et al., 2016), nutrition and milk production (Abdalla et al., 2017), placental homeostasis and uterine environment (Farin et al., 2006; Pohler et al., 2016), embryo-lethal genetic mutations (Pohler et al., 2016), and unbalanced immunological factors at the maternal-fetal interface (Bidarimath and Tayade, 2017).

The global livestock industry depends on the successful use of reproductive techniques to increase the productivity of herds, especially of animals with superior genetics. One of the most useful tools for this purpose is Artificial Insemination (AI), which allows for the maximization of reproductive performance (Baruselli et al., 2012). Currently, it is recognized that Fixed-Time Artificial Insemination (FTAI) is an even more efficient tool to increase the reproductive performance of beef cattle (Baruselli et al., 2019). The efficacy of AI or FTAI procedures can be measured through the pregnancy rate. Therefore, why not associate eGAL with FTAI protocols.

In this context, the prevention of embryonic loss is still a challenge, given the complexity of mechanisms involved in the development and maintenance of pregnancy. Thus, new technologies have been developed, based on knowledge of the cattle’s reproductive physiology, to achieve better reproductive results and reduce production costs, for instance, the hypothesis of using eGAL-1 as presented herein.

During pregnancy, uterine vascularization undergoes dramatic changes with a remodeling of existing vessels and the formation of a new network through angiogenesis that is stimulated (Cross et al., 2002) to provide an adequate supply of oxygen and nutrients for the developing embryo. At the same time, at the embryonic implantation site, the placentation process depends on a complex interaction between invasive trophoblasts and maternal immune cells, involving periods controlled by the development of branched angiogenesis, trophoblast differentiation, and syncytia formation. Interruption/alteration of this pattern of placental development can directly affect its function and result in pregnancy loss in humans (Blois et al., 2019). There are important differences in the degree of invasiveness of blood vessels during placentation of primate and rodent species (greater invasiveness) and ruminants (less invasiveness) and there is still no way to prove that in ruminants, GAL-1 acts on angiogenesis, however it is believed to have an important role in modulating maternal recognition.

Embryos generated by AI procedures consist of 50% genetic material from sperm deposited in the uterus. According to Hide and Schust (2016), alloantigen’s derived from parents or partners are present in the maternal environment at various times during the reproductive process. Embryos originating from embryo transfer procedures, when inovulated in the recipient’s uterus, have a greater chance of maternal-fetal incompatibility, as it is an embryo composed of 100% different genetic material. They are first exposed in semen deposition (either in the natural mating or AI conception) and continue during implantation and placentation. Changes in maternal immune responses allow fertilization and survival/development of the semiallogeneic conceptus until delivery. This immunomodulation must be balanced and continuous, responding appropriately not only to situations of invasion of commensal pathogens in the uterus, cell or tissue damage and any tendency to malignant transformation, as well as against the paternal alloantigen of the conceptus, making it “strange “ to the uterine environment and consequently attacked by the maternal immune system. The absence of this maternal-fetal tolerance or modulation of maternal immunity (the process by which the maternal organism recognizes/accepts the fetus without its immune system attacking it) can be considered as the main cause of embryonic loss (Hyde and Schust, 2016).

Thus, a complex network of metabolic, immunological, and endocrine interactions is activated during the gestational process. Such interactions are necessary to maintain pregnancy and occur at the maternal-fetal interface (on both sides), including cell signaling pathways related to differentiation and growth, vascular development, and immune regulation.

Complex immunoregulatory mechanisms at the maternal-fetal interface must be balanced to activate maternal tolerance against fetal alloantigens and protection against infections and inflammation (Hyde and Schust, 2016). Many mediators are involved in these mechanisms, and galectins, including GAL-1, play a key role, generating great interest in the field of reproductive medicine due to their unique ability to modulate various processes of gestational development and their potential use as biomarkers for gestational disorders (Blois et al., 2019). These facts support the hypothesis for testing the effect of eGAL-1 administration in increasing the pregnancy rate in cattle production.

The present work explored the potential of the administration of exogenous GAL-1 (eGAL-1) to increase the pregnancy rate in beef cattle, combining the administration of a single dose of eGAL-1 with the FTAI technique, and the results represented a significant increase when using this tool. As mentioned in the literature, this experiment remarkably elucidates the effect of GAL-1 on reproductive physiology, positively impacting the development of pregnancy, and consequently, under the item reproductive efficiency in production farms.

## Material and Methods

### Study design

This study analyzed the increase in the pregnancy rate with the administration of a dose of exogenous GAL-1 (eGAL-1), combined with the technique of fixed-time artificial insemination (FTAI) in beef cattle. An effective dose of eGAL-1 means 1 (one) dose of Tolerana® (Inprenha Biotecnologia) whose administration is “extra” but similar to the application of a dose of semen, during the FTAI procedure. One effective dose of eGAL-1 contains 200±10µg of recombinant protein (GAL-1), diluted in 200µL of sterile PBS 1X pH 7.0 buffer solution (Phosphate Buffered Saline with kanamycin sulfate) present in a 0.25mL French-style straw. The definition of what represents an effective dose of eGAL-1 was established in previous experiments (unpublished data), where 5 different doses were tested, always considering use in the same presentation and administration model cited in the present experiment.

Recombinant GAL-1 was obtained through the construction of a heterologous expression vector containing the gene (pET-29a(+)+lgals-1 gene) and purification to obtain active protein, sterile, in its alkylated form and free of endotoxins. The Tolerana® a veterinary product is duly registered with MAPA [Ministry of Agriculture, Livestock and Food Supply] (MAPA/SP 000104-0.000001), and has intellectual property protection, with deposits at INPI and PCT (several countries), both in partnership with the Faculty of Pharmaceutical Sciences of Ribeirão Preto – University of São Paulo, Brazil.

The verification of the effectiveness of GAL-1 was performed by comparing the pregnancy rate in bovine females combining the administration of semen and eGAL-1 (Treated Group - TG) versus the administration of only one dose of semen (Control Group) in the procedure of IA. Thus, in TG, the dose of eGAL-1 is deposited in the lumen of the uterus after the deposition of the semen dose, therefore, there are two procedures for passing the applicators through the cervix. Pregnancy rates for each group (TG and CG) were determined by ultrasound diagnosis (between 28 and 3 days after the FTAI procedure) and submitted to statistical analysis.

### GALECTIN-1 (GAL-1) production and purification

GAL-1 can be obtained from mammalian genomes (from species such as human, bovine, ovine, caprine, equine and porcine) through heterologous expression systems, in the form of active protein, sterile, alkylated, and free of endotoxins. The method for obtaining recombinant Galectin -1 is determined by the manufacturing process of Tolerana® (Inprenha Biotecnologia®) and involves the following steps: (i) obtaining a crude extract of bacteria cultivated to express Galectin-1; (ii) purification of Galectin-1; (iii) preservation of the lectin activity of Galectin-1 by alkylation; (iv) removal of bacterial endotoxin (LPS) from the alkylated Galectin-1 solutions; (v) adjustment of protein concentration; (vi) filling and (vii) quality control. Among the possibilities disclosed in the literature for the upstream and downstream steps, Galectin-1 was produced based on the following procedures, including particularities of the manufacturer’s process. *Subcloning of Gal-1 into pET-29a(+) expression vector*

*Gal-1* Consensus Coding Sequence (CCDS) CCDS13954.1 (length 408nt) was synthesized and subcloned, with juxtaposed insertion of the desired sequence, immediately after the RBS Ribosome binding site sequence of a pET-29a (+) expression vector cut in NdeI / HindIII (GenScript®). This construct was then used for competent transformation of the Rosetta strain of Escherichia coli, maintained in a cell bank.

### Bacterial culture, expression, and Lysis

Aliquots of *E. coli* strains transformed with the insertion of the vector containing the GAL-1 gene (pET-29a(+)+lgals-1 gene) were grown in systems with LB Broth Base medium containing kanamycin sulfate until obtaining optimal bacterial growth rate, demonstrated by optical density. Induction of expression is done with the addition of Isopropyl-D-Thiogalactopyranoside (Sigma-Aldrich) to the culture. After the induced growth period, the bacterial suspension is retained by microfiltration on a Hollow Fiber membrane (0.22µm, Cytiva) and centrifuged at 5000 g for 15-20 minutes at 4°C, always with the supernatant being discarded and the “ bacterial crude = pellet”, which were then subjected to bacterial lysis.

For Bacterial lysis, the crude or bacterial pellet was resuspended in Phosphate Saline Lysis buffer (1X PBS - 136.8 mM NaCl, 2.7mM KCl, 6.4 mM Na2HPO4, 0.9 mM KH2PO4, pH 7.4), containing 14 mM Mercaptoethanol, protease inhibitor EDTA-free, lysozyme-1, RNAse A-Type 3A, and DNAse I Type IV-10. All components are Sigma-Aldrich. The pellet diluted in Lysis buffer (Chemical Lysis) was subjected to constant homogenization for 70 minutes and then sonicated for 3 cycles of 15 seconds each in a Vibra-Cells Sonicator, Sonics (Mechanical Lysis), with intervals of 20 seconds between each cycle. The bacterial lysate was then clarified by centrifugation at 7,000 g for 20 minutes at 4°C and filtered through a 1.0 µm filter (Whatman) with the aid of a peristaltic pump (maximum pressure of 4 BAR).

### Purification steps

After the Chemical and Mechanical Lysis process, the lysate was submitted to 3 steps of purification by chromatography in an AKTA Protein Purification System (Cytiva) to obtain a buffered protein solution containing only Galectin-1.

The first step is based on affinity chromatography on agarose-lactose columns (Sigma-Aldrich), previously equilibrated with equilibration buffer (1XPBS, 14mM 2-ME, pH 7.4). After injection of the protein solution, the affinity column “binders” were washed and eluted with elution buffer (1X PBS containing lactose and 2-ME pH 7.4). The protein peak was collected and 20μM of iodoacetamide (Sigma-Aldrich; I1149) was added to the solution, keeping it under incubation at 4°C, protected from light, overnight. After this incubation, the solution was subjected to “size exclusion” chromatography (Sephadex G-25, Cytiva) to remove the free salts of iodoacetamide and lactose. The last chromatographic step was the removal of bacterial endotoxins (LPS). To this end, the preparations were subjected to chromatography using LPS affinity resin (PIERCE High-Capacity Endotoxin Removal Resin column - Thermo Scientific). After all the chromatographic steps, the protein concentration was determined by spectrometry (Abs 280nm) and expressed in milligrams of protein per milliliter (mg/mL) and were submitted to sterilizing filtration (0.22 μm PES membrane).

Purified protein batches were submitted to the last stage of industrialization only if they reached compliance with the quality standard predetermined by the company, including protein concentration, microbiological status, protein bioactivity (Hemagglutination test), molecular weight analysis by SDS-PAGE, and SEC (size exclusion chromatography), protein secondary structure analysis (Circular Dichroism Analysis), aggregate detection and molecular size by DLS (Dynamic Light Scattering) analysis and endotoxin quantification (LPS). Protein identity was confirmed by LCMS (Liquid Chromatography Mass Spectrometry) and nucleotide sequence confirmation of human galectin-1 cDNA - galectin-1 [Homo sapiens] Consensus Coding Sequence (CCDS) CCDS13954.1 (https://www.ncbi.nlm.nih.gov/projects/CCDS/CcdsBrowse.cgi?REQUEST=ALLFIELDS&DATA=CCDS13954.1&ORGANISM=0&BUILDS=CURRENTBUILDS).

One dose of eGAL-1 translates to 200 ± 10 µg of purified protein diluted in 200 µL of Sterile Phosphate Buffer solution (1X PBS pH 7.0) containing 50 µg/mL of kanamycin sulfate. The commercial presentation of eGAL-1 (Tolerana) is in a paper box containing 50 straws stored in vacuum-sealed plastic containers and kept at 5 ± 3°C until the moment of use. The material was transported in isothermal boxes containing hard ice (5 ± 3°C).

### Field experiment

#### Location

The experiments were conducted in 17 commercial beef cattle farms located in different Brazilian municipalities (Campo Grande - MS; Naviraí - MS; Água Clara - MS; Formoso do Araguaia - TO; Gurupi - TO; Paragominas - PA; Uberaba - MG, Uberaba – MG; Pedregulho – SP; São Gotardo – MG; Prata – MG; Água Clara – MS and Cuiabá - MT). It should be noted that the farms selected to participate in this experiment were farms that have a history of working with FTAI procedures for at least 2 years.

#### Animals

The experiments were carried out on female bovine animals (cows conventionally managed as dams and not intended for slaughter) managed in extensive beef cattle rearing systems. The dams were kept in an extensive rearing system, under native and/or cultivated pasture, with mineral supplementation. All cows underwent an FTAI procedure. It should be noted that only cows diagnosed as empty (by ultrasonography, 28 to 35 days after the first service - 1^st^ FTAI) were worked on in a second FTAI protocol. Fifteen days from the 2^nd.^ FTAI, bulls were introduced for transfer with natural breeding. It is important to remember that the experiment and the statistical model considered only 1^st^ service results to ascertain the effectiveness of the dose of eGAL-1. Prophylactic management with annual vaccinations against BHV-1, BVDV, and BL (dose and booster) of cows before the start of the breeding season was implemented in some farms.

In total, 3469 beef cows (Nellore and crossbred dams) were considered in the statistical model, which were divided into 2 treatment groups (TG and CG) and for statistical analysis divided into 107 contemporary groups (YG) as described below. The experiment was designed with 4730 dams at the time of insemination, distributed equally (same number of cows in each group n= 2365) and randomly (without the previous choice of the TG or CG group that would be part of). However, 1261 cows were excluded from the experiment, because different reasons, including (i) dams did not maintain BSC between 3.5 and 2.5; the dams’ body condition score (BSC) was observed in two situations – at the time of the FTAI and the day before the pregnancy diagnosis. Only dams that maintained a BSC between 3.5 and 2.5 in the 02 situations mentioned above were approved to participate in the statistical analysis (ii) dams died during this interval; (iii) became ill during this interval (e. g., hoof, mastitis, diarrhea, pneumonia); (iv) those who had problems with the synchronization protocol (e. g. loss of CIDR); (v) who changed management lot); and for these reasons, it is noted that in some farms the number of dams mentioned in table 03 differs between the TG and CG groups. The numerical decompensation between groups was corrected in the construction of contemporary groups (YG), as described in the item below (Contemporary Groups and Statistical Analysis). An important detail is that the dams were submitted to BSC classification before the pregnancy diagnosis.

The criteria for defining the BSC used were based on the descriptions by Machado et al.(2008), who empirically determined the separation of dams into 5 BS classifications: 1 (cachectical): complete visualization of the ribs, exposures of ileum bones and ischium, and pronounced muscle atrophy (skin and bones apparent); 2 (thin): very prominent bones with visible dorsal, iliac and ischial processes; 3 (great): light muscle coverage and no fat accumulation; 4 (fat): good muscle coverage and fat deposition at tail insertion; 5 (obese): all body angles covered, including protruding skeletal parts and overall animal appearance.

The experiment considered 3 different animal categories in the work lots – heifers, multiparous and primiparous. Multiparous and Primiparous cows had calves on their feet, at 60 to 100 days of lactation. These categories defined differences in the estrus synchronization protocols used for the categories.

### FTAI and eGAL-1 administration

The breeding cows were kept in management batches on the farms. Each batch was submitted to FTAI after estrus synchronization protocols were performed. These synchronization protocols were decided by each farm, as described in Table 01. We did not interfere in these protocols and within each batch, there were no changes in the protocols. No imposition was imposed on participating farms in the choice of estrus synchronization protocols. Other decisions by the participating farms were (i) regarding the selection of the “bull” (semen doses) selected for use in the FTAI procedure and (ii) the choice and training of the inseminator who would inseminate each batch of breeding cows. A total of 46 bulls were used, selected by the partner farm and the semen doses of each bull, distributed in the Treated (TG) and Control (CG) groups. In total, 23 inseminators participated in the experiment carried out on these 17 farms. There was no previous selection for the dam to receive the dose of eGAL-1 during the AI procedure. If the first dam that entered the containment trunk received the dose, the second did not receive it, thus continuing until the end of the insemination of the batch in question.

**Table 01.** Farm (in letter codes), category of cows (H = heifers, P = primiparous or M = multiparous), days of estrus synchronization protocols (D0= day zero, D7 = day seven, D8= da eight, D9 = day nine, TAI day = day of the AI procedure (CG) and eGAL-1 and respective hormones applied, with amounts administrated, according to the estrus synchronization protocol adopted by the partner farm.

| Farms Codes | Category | D0 | D7 | D8 | D9 | TAI day |
| --- | --- | --- | --- | --- | --- | --- |
| A | P | EB<br>2.0mL | - | - | PGF<sub>2α</sub> 2.0mL | D11 |
|  |  |  |  |  | ECP 1.0mL<br>eCG 1.5 mL |  |
| K | M | EB<br>2.0mL | - | - | PGF<sub>2α</sub> 2.0mL<br>ECP 1.0mL<br>eCG 1.5 mL | D11 |
| B | M | EB<br>2.0mL | - | PGF<sub>2α</sub> 1.5mL<br>ECP 0.5mL<br>eCG 1.5 mL | - | D10 |
| C | M | EB<br>2.0mL | PGF<sub>2α</sub><br>2.0mL | - | ECP 0.5mL<br>eCG 1.5 mL | D11 |
| F<br>J<br>L<br>K<br>M<br>N<br>O | H | EB<br>2.0mL | - | PGF<sub>2α</sub> 2.0mL<br>ECP 1.0mL<br>eCG 0.5 mL | - | D10 |
| G<br>H<br>I<br>J<br>L<br>M<br>N<br>O<br>P<br>Q<br>R<br>T | M | EB<br>2.0mL | - | PGF<sub>2α</sub> 2.0mL<br>ECP 1.0mL<br>eCG 1.5 mL | - | D10 |
| J | P | EB<br>2.0mL | - | PGF<sub>2α</sub> 2.0mL<br>ECP 1.0mL<br>eCG 1.5 mL | - | D10 |

EB- Estradiol Benzoate. PGF<sub>2α</sub> -Prostaglandin F<sub>2</sub>alpha. EC- Estradiol Cypionate. eCG
– Equine Chorionic Gonadotropin

The procedure in the treated group TG was to inseminate the breeding cows using a conventional semen applicator, followed by the administration of the eGAL-1 dose using a second applicator (identical to the semen), which represents that breeding cows in the treated group were at disadvantage compared to the CG, as they had 2 events of the transgression of the cervical rings. The deposition of the eGAL-1 dose in the uterine lumen was performed as the second insemination where after removal of the semen applicator, a second applicator mounted with a straw containing the protein dose was re-introduced as shown in Fig 1.

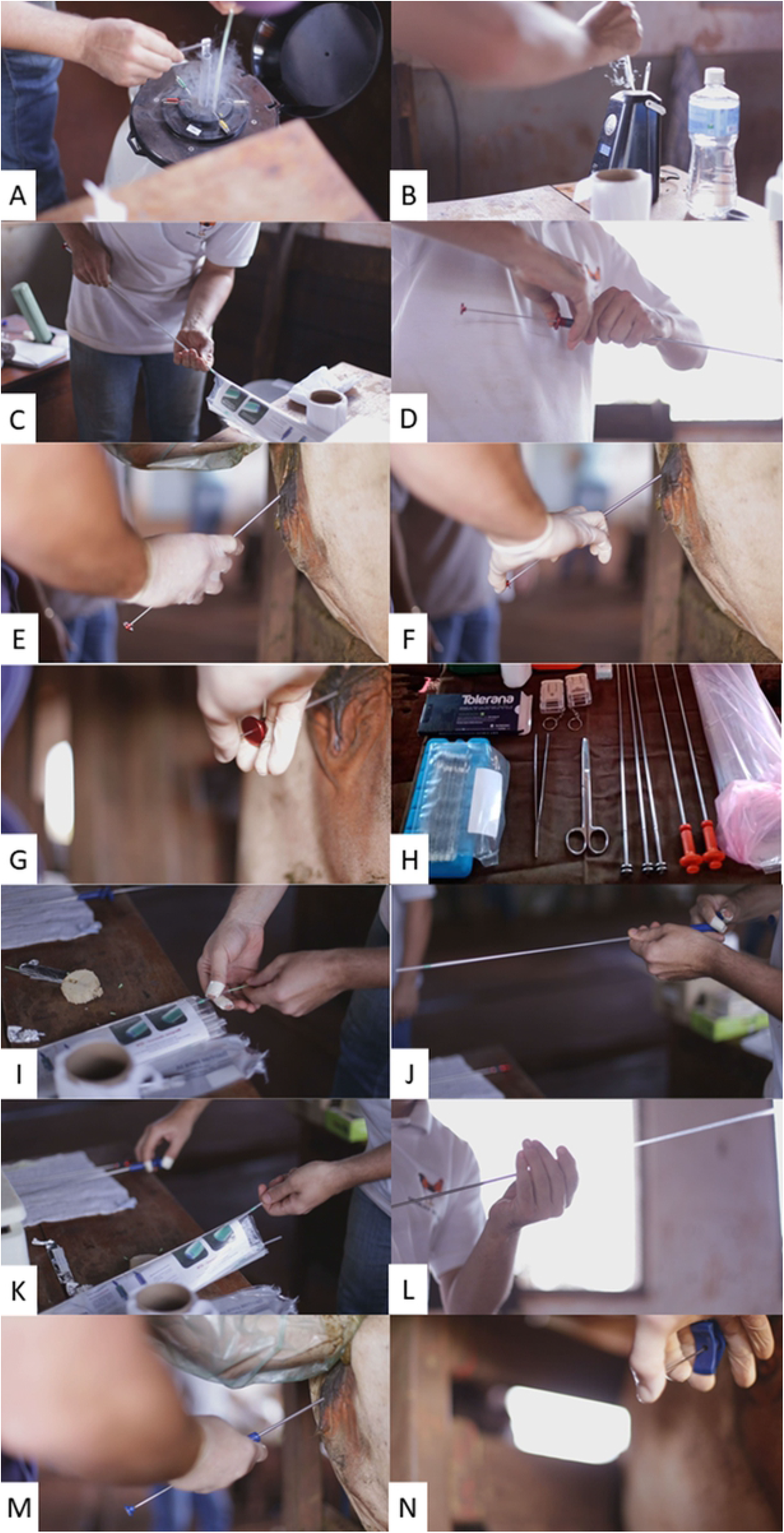

As a procedure in group CG, the females were inseminated according to the standard procedure, the same one recommended by (Brazilian Association of Artificial Insemination, 2018), with a single dose of semen (which represents 1 applicator being passed through the cervical rings). The time spent for the insemination procedure in the females of the CG and TG groups was considered a point of attention to the method. In this experimental model, we worked with the prerogative that dams belonging to the CG were at an advantage compared to the TG, as they received only “one act” to transverse the cervical rings during the procedure, and that it is also known that this “act “ can negatively affect the pregnancy rate.

### Obtainment of the pregnancy rate in the groups

The experimental results of pregnancy rates obtained in the breeding cows of the TG and CG groups obtained by ultrasonography (28 to 35 days after FTAI), and the increase in the rate obtained using the eGAL-1 was calculated based on statistical methodology considering dams, which maintained a BSC between 3.5 and 2.5, divided into 2 experimental groups (TG and CG) compared within the same contemporary groups formed, as described below. The diagnosis was performed by a technician with experience in ultrasonography and without knowledge of the division of dams into TG and CG groups.

### Contemporary groups and statistical analysis

To define and compare the pregnancy rates between breeding cows inseminated with and without Tolerana®, they were grouped into contemporary groups (YG). Each contemporary YG group was composed of dams inseminated by the same inseminator (identified by a letter code), belonging to the same farm (identified by a letter code), of the same animal category (which codes were used to facilitate separation (M = multiparous, P = primiparous or H = heifers), from the same management group (identified by the FTAI date + Farm code + management lot code); inseminated with the same semen batch (identified by the name of the Bull) and being of the same breed (N = Nellore and CB = crossbreed) A minimum number of at least 5 dams was considered to form a YG or those groups that did not show variation in the pregnancy rate (100 or 0%) were also discarded. To perform the analysis, the Generalized Linear Model (GLM) was applied, with the GENMOD procedure of the SAS (version 9.3), assuming a binomial distribution (pregnant or not pregnant) with residual effect and a logarithmic function (PROBIT). The model included the fixed effect of YG and treatment (dose = 0 of eGAL-1 in the CG and dose = 200 that means 200 ± 10 µg of eGAL-1 in the TG).

Altogether, 107 YG were formed with the arrangement of these 3469 breeding cows, distributed among the 3 animal categories, inseminated with doses of semen from 46 bulls, by 23 inseminators and distributed among the farms and management batches. Remind that 1,261 dams were excluded to the statistical analyses as mentioned previously.

The PROC GENMOD is modeling the probability that Pre=‘2’ = pregnant, using Information (Prm) and Effect parameters, where Prm1 = Interception; Prm2 = Dose0; Prm3 =Dose200; Prm4 = YG#1; Prm5 = YG#2; …..Prm110 = YG#107. Dose200 means administration of eGAL-1 in TG dams.

The pregnancy rate obtained in these 107 YG was determined considering the diagnosis of pregnancy by ultrasound between 28 and 35 days after the FTAI procedure.

### Ethical Statement

This study complied with the ethical requirements for the use of animals in experiments and was approved by the CEUA/USP, protocol number 11.1.95.53.5.

## Results

The administration of the recommended dose of eGAL-1 for the FTAI procedure took about 10 ± 5 seconds longer than a conventional procedure. The probability of positive pregnancy in the CG group was 49.4% while in the TG group it was 58.08% (p<0.0001), as detailed in Table 2. The mean obtained with Dose0 = 0.491, equivalent to 49.41% of probability of obtaining a positive pregnancy in the CG, while the average obtained with Dose200 = 0.5808, equivalent to 58.08% probability of obtaining a positive pregnancy in the TG, which represents 8.68% difference between the treatment groups, when compared within of each YG.

**Table 2.**
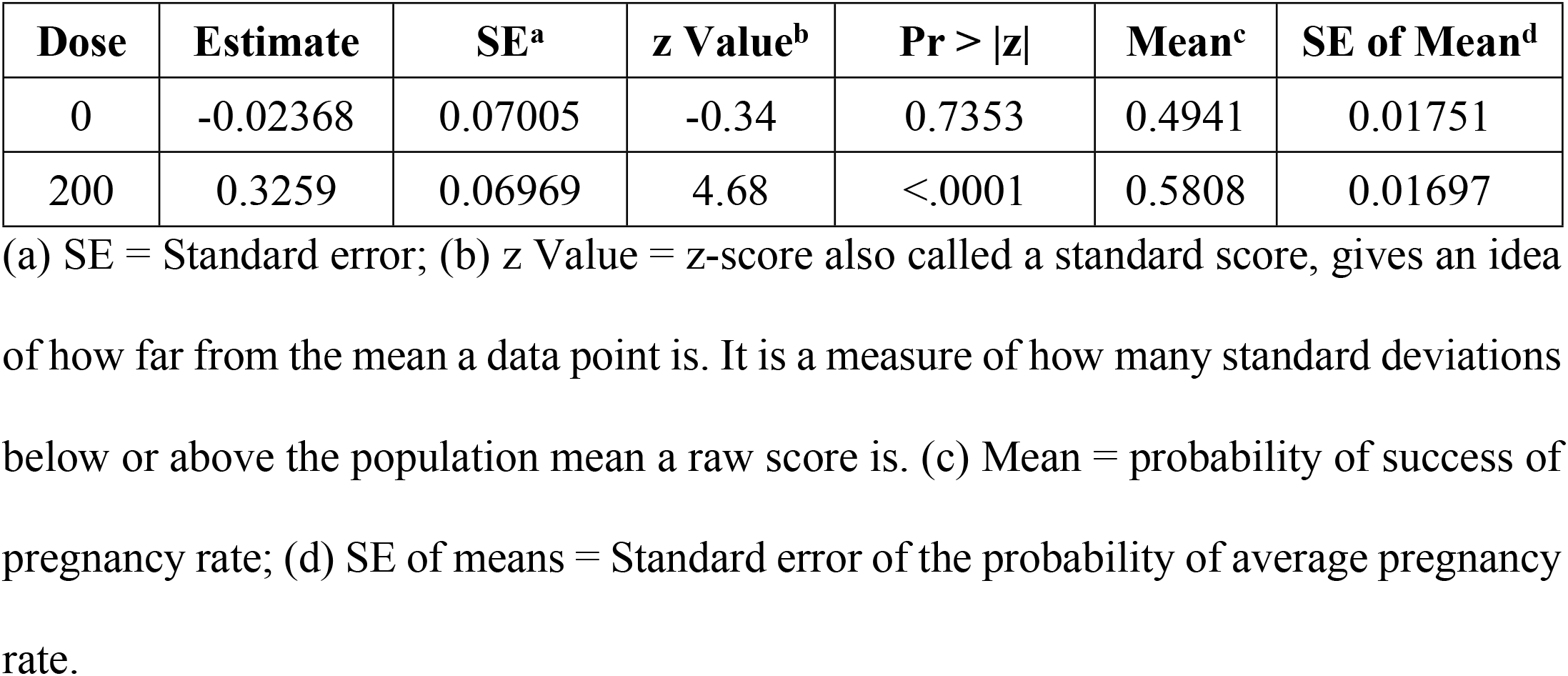
Dose Least Squares Means, using Generalized Linear Model (GLM), with GENMOD procedure, under binomial distribution (pregnant/ not pregnant) and in logarithmic function (PROBIT), by source as “dose of eGAL-1”, that means dose 0 = GC and dose 200 = TG, using [SAS] software, Version [6.9].

The “YGs effects”, under binomial distribution (“pregnant” and “not pregnant”), did not present statistical significance (p = 0.1787), perhaps because the variables that made up the construction of the YG greatly interfere in the pregnancy rate.

Table 03 describes the simple average obtained in each group (TG and CG) in the different farms, within each animal category, and for each inseminator who performed the procedures. Based on the comparison between the simple means of the CG (48.58%) and the TG (58.34%), a 9.76 percentage point difference was obtained between the groups. We are aware that the pregnancy rate can be interfered with by several factors or variables, going well beyond “just the location of the farm”, so we proposed to discuss a discussion based on results based on this proposed statistical model, which considers “product dose-effect” within contemporary groups, grouping all impacting variables for “pregnancy rate” within each of the YG created. There were so many variables that 107 YG were created in the established statistical model. Importantly, among the calves born in this experiment, more than 900 of them conceived and gestated in the uterus that had contact with the eGAL-1 protein (dams belonging to the TG group) during the FTAI procedure, no congenital defect or stillbirths were observed.

**Table 03.**
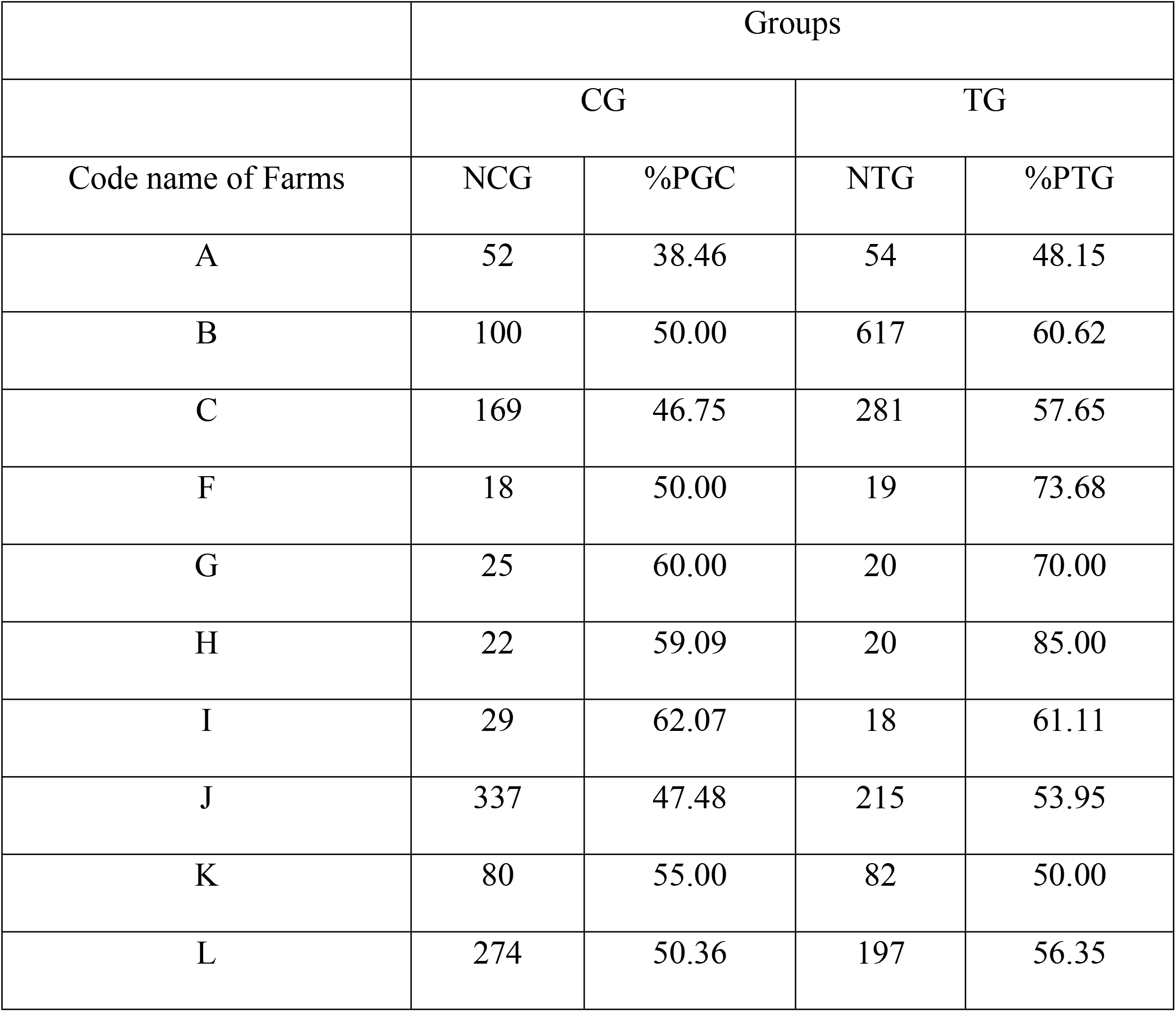
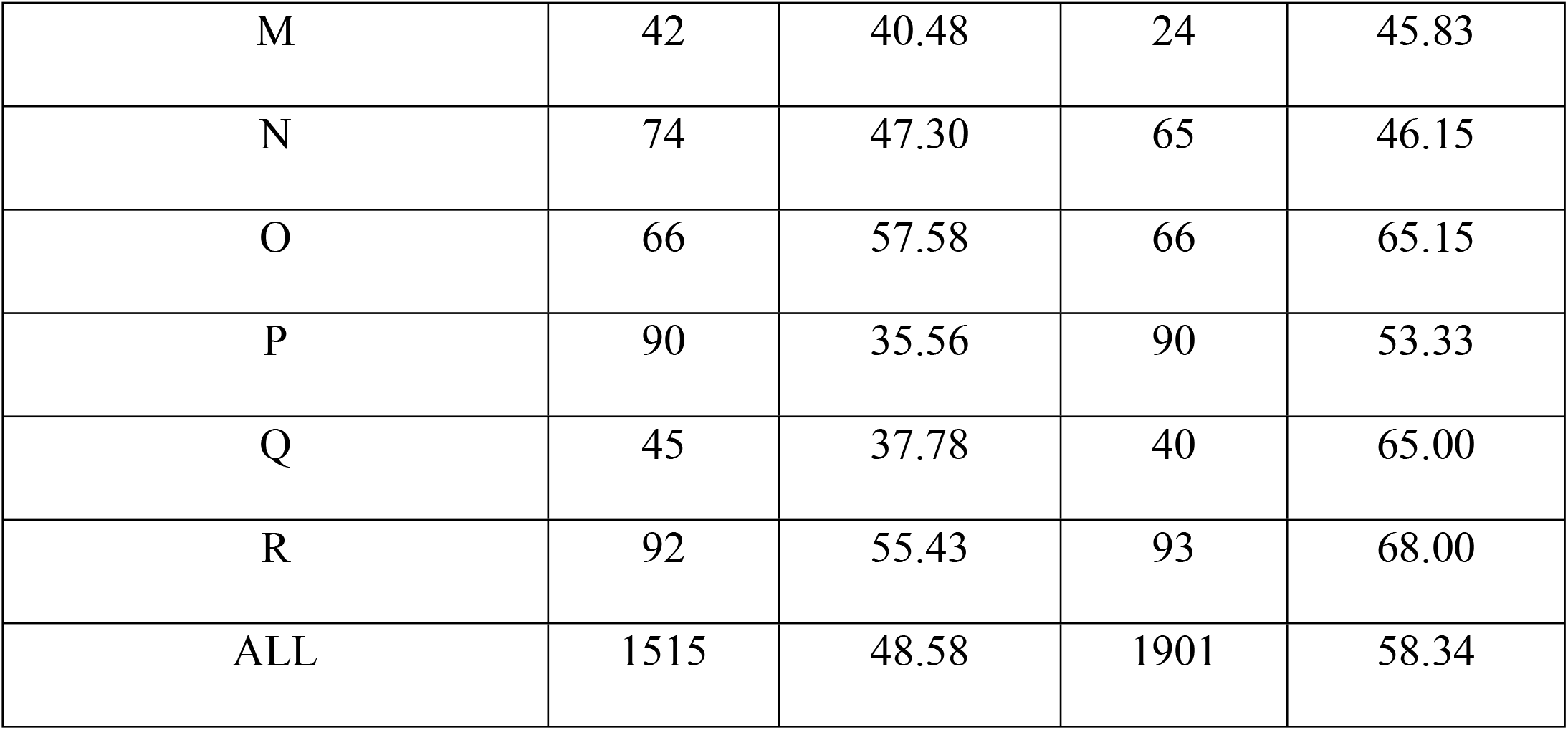
Average of pregnancy rate (%P) and number of cows (N) on the Control Group (CG) and Treated Group (TG) by each farm (identified by code name) and by all farms. Font in bolt format indicate higher %P in TG.

|  | Groups |  |  |  |
| --- | --- | --- | --- | --- |
|  | CG |  | TG |  |
| Code name of Farms | NCG | %PGC | NTG | %PTG |
| A | 52 | 38.46 | 54 | 48.15 |
| B | 100 | 50.00 | 617 | 60.62 |
| C | 169 | 46.75 | 281 | 57.65 |
| F | 18 | 50.00 | 19 | 73.68 |
| G | 25 | 60.00 | 20 | 70.00 |
| H | 22 | 59.09 | 20 | 85.00 |
| I | 29 | 62.07 | 18 | 61.11 |
| J | 337 | 47.48 | 215 | 53.95 |
| K | 80 | 55.00 | 82 | 50.00 |
| L | 274 | 50.36 | 197 | 56.35 |
| M | 42 | 40.48 | 24 | 45.83 |
| N | 74 | 47.30 | 65 | 46.15 |
| O | 66 | 57.58 | 66 | 65.15 |
| P | 90 | 35.56 | 90 | 53.33 |
| Q | 45 | 37.78 | 40 | 65.00 |
| R | 92 | 55.43 | 93 | 68.00 |
| ALL | 1515 | 48.58 | 1901 | 58.34 |

## Discussion

### Efficacy of exogenous Galectin-1 on the increase of pregnancy rate

The aim of this study was to explore the potential effect of eGAL-1 in increasing the pregnancy rate in beef cattle, combining the FTAI technique, the administration of a single dose of eGAL-1, measuring the difference in efficacy obtained, through the designed statistical model (comparison of the pregnancy rate between TG and CG within the formed YG), and thus it was observed that the probability of the pregnancy rate was 58.08% in TG and 49.4% in CG (p<0.0001), demonstrating a positive efficacy of eGAL-1 on the pregnancy rate (Table 2).

YG fixed several factors that can interfere with the pregnancy rate in beef cattle, keeping only the variable dose of eGAL-1 as an extra variable in the statistical model. YG considered the grouping of breeding cows from both groups (TG and CG) equally distributed (before dams’ exclusion) within the same batch, within the same farm, within the same category, inseminated with doses from the same bull and by the same inseminator. Particulars of sanitary and nutritional management are also being considered in the YG grouping (batch). The negative effect on the pregnancy rate (or increased probability of pregnancy loss) due to nutritional deficiency was controlled in the experiment, excluding dams that did not maintain their body score, as previously mentioned.

Thus, the pregnancy rate obtained on TG was different, (in this case, 8.68% higher), only because of the eGAL-1 administration (p<0.0001). Remembering yet, the YG effect, under binomial distribution (pregnant or not pregnant), was not statistically significant (p = 17.87). However, in the scenery with 3469 dams, 3 animal categories, 46 different bulls, 23 inseminators, several batches, and 2 treatment groups, it is reasonable to consider the YG effect as a biological effect. The recommendation to use a dose of eGAL-1 during a FTAI procedure was reasonable in the beef cattle routine. On average, the whole procedure, when we administrate de eGAL-1, spent only 10 ± 5 seconds more than the conventional procedure - ten seconds as a price to get 8.68% more chance to pregnant a dam, is reasonable in the animal production systems.

The form of eGAL-1 administration is not foolproof to improve the pregnancy rate in beef cattle but showed that can help. eGAL-1 means Tolerana® administration and it is an innovative technology and its efficacy experimentation model was executed between the manufacturing company and partner farms. However, as in any product development process, we go through stages and technological challenges, such as (1) the ideal dosage for different cows categories, (2) the ideal eluent for maintaining the stability of the active protein in the product, (3) ideal packaging to facilitate the procedure and to not harm the product, (4) determination of the best form and moment of application, (5) the application procedure definition, (6) the interference of synchronization protocols, among others. In this scenario, before obtaining this 8.68% higher pregnancy rate achieved in this experiment, using eGAL-1 administration, some procedures might not work well.

Also, there were technological challenges in manufacturing the active ingredient - a recombinant protein that has intrinsic particularities to the molecular structure, and it can suffer the methodology influence and the manufacturing processes. It should be noted that, like any biotechnological product, “product = process”, while the product had no commercial registration, optimizations were performed in the process, aiming for greater yield and scale-up manufacturing. In some situations, with certain partners, such changes had a negative impact on the stability and effectiveness of the technology, resulting in no or small increase in the pregnancy rate. However, the manufacturing process is currently consolidated and robust on homogeneity, stability, and efficiency. In fact, during the experimental phase of technology development (Tolerana®), it should be noted that it was used more than 12 thousand times in AI procedures of bovine dams, and there were no undesired biological effects. There were no reports of discomfort, pain, and irritation with the administration of the product, except for those already known in artificial insemination procedures. It is also important to note that stillbirths, malformations, and/or neonatal complications were not verified. There are no reports of intoxication in humans using GAL-1 as an active ingredient in drugs. In the current literature, galectins (soluble in blood serum or expressed in tissues) have generally been used as biological markers of several pathological events (He et al. 2017; Vergetaki et al., 2014; Vasta, 2012), directing treatments, but still in experimental stages.

Another important note about interference on results falls on semen doses. The doses of semen used in this experiment were a “farm decision”, even that we observed 46 different bulls, used. eGAL-1 is a protein and is feasible occur an interaction between it and some components of the semen extender, including lactose, a carbohydrate used in some recipes of semen extender that present a high interaction with galectin 1. If the protein binds with the semen extender lactose, it will be interacting with the endometrium at the time of administration of eGAl-1? This is a hypothesis does not respond by the authors yet. Difficult to know every semen extender present on semen doses used. Centers that industrialized semen doses do not share this information easily. For this reason, YG was considered as statistical analysis.

The efficacy of the technology should also be observed carefully. If it has not been performed consistently, considering the experimental and statistical model (as described in item *Methods* - *Contemporany groups and statistical analysis*), the effect of effectiveness can be masked. Thus, when used correctly, the results are quite promising. The indication of Tolerana® is NOT for the treatment of infertile nor sick animals, but it is indicated as a health catalyst of animal fertility, being, therefore, a tool to increase the reproductive/productive efficacy, impacting economically on the productive cattle chain.

### The economic impact of reproductive efficiency must be calculated and/or considered

In the beef cow-calf system, the number of calves on the final breeding season, number of days up to the time of a new pregnancy (feeding cost versus open days), and cow reposition taxes must be considered the principal goals for a bio-economic evaluation. Moreira (2019) said that it is possible to reduce the investment cost of the FTAI program by 10% for each percentage point added to the pregnancy rate. If true, using eGAL-1 administration on FTAI procedure the consumer can reduce the cost with the FTAI program per cow considerably and, consequently, can also reduce the cost of calves’ birth. The cost benefits of eGAL-1 are interesting, considering that a dose of Tolerana® means 10% plus on the cost of the FTAI program per dam. More details of the economic impact of using eGAL-1 on the pregnancy rates rise in FTAI programs will be discussed in another paper.

The data presented in this study corroborate with several authors, as described below, and support the innovative hypothesis and the new product presented.

### Why galectin-1 could improve the pregnancy rate?

Galectins are a family of evolutionarily conserved proteins distributed from lower invertebrates to mammals (Cummings & Liu, 2009; Modenutti et al., 2019). Thus, the efficacy of recombinant human Gal-1 under assisted reproduction procedures was evaluated, besides bovine females (this work), ovine, and equine (unpublished data). All evaluations presented in species different from bovine promising results only with dosage adjustment because of the area (cm^2^) of lumen uterus of each species.

From the mid-1970s onwards, several findings of animal lectins and β-galactoside ligands have been described. Barondes et al. (1994a) proposed the creation of the galectin family to group these proteins. “Electrolectin” was the first member of that family, isolated from tissues of electric fish (Teichberg et al., 1975). Even the first β-galactoside-binding lectins derived from mammals were described and were later defined as galectin-1 and 3 (Barondes et al., 1994b).

There is enormous structural diversity of glycoconjugates in living beings, which can be associated with a significant biological diversity since these glycostructures can encode various biological information decoded by lectins (Sharon and Lis, 1989). Therefore, the carbohydrates recognition by lectins is a biochemical phenomenon associated with several physiological and/or pathological processes such as cell fertilization, embryogenesis, migration, proliferation, and differentiation, immune defense, infection by microorganisms, and cancer (Sharon & Lis, 1986; Sharon & Lis, 1989; Santos-de-Oliveira et al., 1994; Dias-Baruffi et al., 2003; Liu & Rabinovich, 2005).

Fifteen mammalian-derived galectins have been described, and all have a CRD with approximately 130 amino acid residues. Galectins were classified into three categories: “proto-type”, chimera, and “tandem repeat-type”, the first being those with a single CRD type and with identical monomers or dimers with CRD associated non-covalently (galectins: 1, 2, 5, 7, 10, 11, 13, 14 and 15). Galectin-3 represents the chimera type, has a CRD, and a non-lectin domain involved in its oligomerization (Cummings & Liu, 2009; Modenutti et al., 2019). Galectins 4, 6, 8, 9, and 12 are known as “Tandem repeat-type” because they have two distinct CDRs joined by a small binding peptide (Rubinstein et al., 2004a).

Most galectins have characteristics of cytoplasmic proteins such as acetylated N-terminal region, non-oxidized (free) sulfhydryl groups, and absence of glycosylation (Rubinstein et al., 2004a and b). However, galectins can be located on the cell surface, extracellular matrix, cytoplasm, and cell nucleus (Rubinstein et al., 2004a). Although these proteins can be detected in the extracellular environment, they do not present signal peptides, being secreted by the cells by a non-classical mechanism and independent of the endoplasmic reticulum and the Golgi complex (Hughes, 1999). Literature data suggest that GAL-1 can be secreted by direct translocation of cytosol through the plasma membrane with the aid of cytosol and membrane factors, as described for fibroblast growth factor-2 (Schäfer et al., 2004; Nickel, 2005).

Since galectins are bivalent, they can promote the intercrossing of glycoconjugates on the cell surface in the extracellular environment and induce signal transduction events by forming clusters of receptors and a mesh (galectin-receptor) on the cell surface (Brewer et al., 2002). In the intracellular environment, the biological events of galectins do not seem to depend on their lectin properties, and they can participate in the processing of RNA and the regulation of cellular homeostasis (Liu et al., 2002; Wang et al., 2004). Interestingly, galectins can exert antagonistic effects depending on these proteins’ location in the intra- or extracellular environment (Yang et al., 1996; Fukumori et al., 2003). Then et al. (2008) showed that LGALS1 has a high degree of structural conservation, dimerization, and binding properties with carbohydrates and integrins (adhesion proteins), suggesting that these properties are conserved among vertebrates and that they maintain a pattern of gene expression among the different types of the placenta (deciduous or not).

Galectins are multifunctional molecules that participate in several biological processes such as adhesion, proliferation, and cell cycle, apoptosis, RNA processing, control of the inflammatory process, and physiological mechanisms of reproduction (Perillo et al., 1995; Liu et al., 2002; Dias-Baruffi et al., 2003; Rubinstein et al., 2004b; Stowell et al., 2007; Ramhorst et al., 2012; Barrientos et al., 2014; Blois et al., 2019).

The galectin’s maternal-fetal tolerance role, both innate and adaptive, is associated with regulating and modulating the embryo elongation events’ immunological responses and adherence to the endometrium. Besides GAL-15 and GAL-1, other galectins can be expressed by the endometrium and the placenta of mammals, presenting essential functions in differentiating the endometrium implanting the blastocyst, and differentiating the trophoblast (Farmer et al., 2008). They also contribute to placentation as they regulate the development, migration, and trophoblastic invasion, essential in early gestational development (Barrientos et al., 2014; Blois et al., 2019, 2007; Freitag et al., 2013). Even, they act in the maternal immunological tolerance mechanism to fetal alloantigen’s, regulating the Natural Killer uterus cells and modulating T cells, which are mainly responsible for cellular immunity (Than et al., 2008).

The endometrial expression of GAL-1 fluctuates during the estrous cycle of different phases because steroidal hormones influence it. GAL-1 has been detected in 3- to 5-day old human embryos, acting on trophoblasts differentiation in the fetus’s placenta and internal cell mass. The interaction of GAL-1 with integrins suggests participation in the extracellular matrix and placentation events, either in the oxygen exchange and/or nutrients or by the vessels formation (angiogenesis), showing that GAL-1 plays a vital role in interface signaling maternal-fetal since it has multiple biological functions (Choe et al., 1997).

Blois et al. (2007) demonstrated high pregnancy loss rates in mice in which the *Lgals1* gene was deficient (knockout mice). When treating deficient mice with recombinant GAL-1, there was a decrease in fetal loss and the restoration of tolerance through several mechanisms, including the induction of tolerogenic dendritic cells, which in turn promoted the expansion of regulatory T cells secreting interleukin-10 (IL-10) *in vivo*. Consequently, the protective effects of GAL-1 have been revoked in mice depleted of regulatory or IL-10 deficient T cells. Thus, they (Blois et al. 2007) demonstrated the fundamental importance of GAL-1 in fetomaternal tolerance and the synergy between GAL-1 and progesterone in maintaining pregnancy.

## Conclusion

This study showed the eGAL-1’s effectiveness in improving the beef cattle pregnancy rate and by administering the recommended dose, the procedure may take 5 to 10 seconds longer than the conventional procedure, however, the statistics show a considerable increase in the pregnancy rate. Considering the “eGAL-1 administration effect” it is possible to improve in 8.68% the chances of pregnancy in an inseminated cow.

## Acknowledgments

Special thanks to Professor Dr. MARCELO DIAS BARUFFI (FFRP / USP) and to all his academicians who collaborated with the initial experiments in the development of the innovative technology. We would also like to thank the inventors of the mentioned patents CAMILLO DEL CISTIA ANDRADE, LÍLIAN CATALDI RODRIGUES, MARCELO DIAS BARUFFI, MARCELO RONCOLETTA, and ERIKA DA SILVA CARVALHO MORANI who ceded/granted the transfer, without reservations, all rights, ownership, action, and interests of the invention to USP (University of Sao Paulo) and Inprenha Biotecnologia. This acknowledgment is extended to USP, co-author of the patent, which transferred to Inprenha Biotecnologia the rights of exploitation of the patent.

## References

1. Abdalla, H., Elghafghuf, A., Elsohaby, I., Nasr, M. A. (2017). Maternal and nonmaternal factors associated with late embryonic and early fetal losses in dairy cows. Theriogenology, 100, 16–23. doi: 10.1016/j.theriogenology.2017.04.005

2. Associação Brasileira de Inseminação Artificial. (2018). Manual de inseminação artificial em bovinos (??rd ed.), ASBIA

3. Barondes, S. H., Castronovo, V., Cooper, D. N. W., Cummings, R. D., Drickamer, K., Felzi, T., et al. (1994a). Galectins: a family of animal beta-galactoside-binding lectins. Cell, 76, 597–598. doi: 10.1016/0092-8674(94)90498-7

4. Barondes, S. H., Cooper, D. N. W., Gitt, M. A., Leffler, H. (1994b). Galectins. Structure and function of a large family of animal lectins. Journal of Biological Chemistry, 269, 20807–20810. doi: 10.1016/S0021-9258(17)31891-4

5. Barrientos, G., Freitag, N., Tirado-González, I., Unverdorben, L., Jeschke, U., Thijssen, V. L. J. L., et al. (2014). Involvement of galectin-1 in reproduction: Past, present and future. Human Reproduction Update, 20:175–193. doi: 10.1093/humupd/dmt040

6. Baruselli, P. S., Sales, J. N. S., Sala, R., Vieira, L. M., Filho, M. F. S. (2012). History, evolution and perspectives of timed artificial insemination programs in Brazil. Animal Reproduction, 9, 139–152.

7. Baruselli, P. S., Catussi, B. L.C., de Abreu, L. A., Elliff, F. M., da Silva, L. G., Batista, E. S., et al. (2019). Evolução e perspectivas da inseminação artificial em bovinos. Revista Brasileira de Reprodução Animal, 43, 308–314. http://cbra.org.br/portal/downloads/publicacoes/rbra/v43/n2/p308-314%20(RB812).pdf

8. Bidarimath, M., Tayade, C. (2017). Pregnancy and spontaneous fetal loss: A pig perspective. Molecular Reproduction and Development, 84(9), 856–869. doi: 10.1002/mrd.22847

9. Blois, S. M., Dveksler, G., Vasta, G. R., Freitag, N., Blanchard, V., Barrientos, G. (2019). Pregnancy galectinology: Insights into a complex network of glycan binding proteins. Frontiers in Immunology, 10, 1166. doi: 10.3389/fimmu.2019.01166

10. Blois, S. M., Ilarregui, J. M., Tometten, M., Garcia, M., Orsal, A. S., Cordo-Russo, R., et al. (2007). A pivotal role for galectin-1 in fetomaternal tolerance. Nature Medicine, 13(12), 1450–1457. doi: 10.1038/nm1680

11. Brewer, C. F., Miceli, M. C., Baum, L. G. (2002). Clusters, bundles, arrays and lattices: novel mechanisms for lectin–saccharide-mediated cellular interactions. Current Opinion in Structural Biology, 12, 616–623. doi: 10.1016/S0959-440X(02)00364-0

12. Cheng, Z., Abudureyimu, A., Oguejiofor, C. F., Ellis, R., Barry, A. T., Chen, X., et al. (2016). BVDV alters uterine prostaglandin production during pregnancy recognition in cows. Reproduction, 151, 605–614. doi: 10.1530/REP-15-0583

13. Choe, Y. S., Shim, C., Choi, D., Lee, C. S., Lee, K. K., Kim, K. (1997). Expression of galectin-1mRNAin the mouse uterus is under the control of ovarian steroids during blastocyst implantation. Molecular Reproduction and Development, 48, 261–266. doi: 10.1002/(SICI)1098-2795(199710)48:2<261::AID-MRD14>3.0.CO;2-0

14. Cobuci, J. A., de Abreu, U. G. P., Torres, R. D. A. (2006). Formação de grupos contemporâneos em bovinos de corte. Embrapa Pantanal-Documentos. ISSN: 1517-1973

15. Cross, J. C., Hemberger, M., Lu, Y., Nozaki, T., Whiteley, K., Masutani, M., et al. (2002). Trophoblast functions, angiogenesis and remodeling of the maternal vasculature in the placenta. Molecular and Cellular Endocrinology, 187, 207–212. doi: 10.1016/s0303-7207(01)00703-1.

16. Cummings, R. D., Liu, F. T. (2009). Galectins. In A. C. Varki, R. D. Cummings, J. D. Esko, H. H. Freeze, P. Stanley, C. R. Bertozzi, G.W. Hart & M. E. Etzler (Eds.), Essentials of Glycobiology (2th ed., chapter 33). Cold Spring Harbor Laboratory Press. PMID: 20301239

17. Dias-Baruffi, M., Zhu, H., Cho, M., Karmakar, S., McEver, R. P., Cummings, R. D. (2003). Dimeric galectin-1 induces surface exposure of phosphatidylserine and phagocytic recognition of leukocytes without inducing apoptosis. Journal of Biological Chemistry, 278, 41282–41293. doi: 10.1074/jbc.M306624200

18. Diskin, M. G., Morris, D. G. (2008). Embryonic and early fetal losses in cattle and other ruminants. Reproduction in Domestic Animals, 43 (Suppl 2), 260–267. doi: 10.1111/j.1439-0531.2008.01171.x

19. Diskin, M., Waters, S., Parr, M., Kenny, D. (2016). Pregnancy losses in cattle: potential for improvement. Reproduction, Fertility and Development, 28, 83–93. doi: 10.1071/RD15366

20. Farin, P. W., Piedrahita, J. A., Farin, C. E. (2006). Errors in development of fetuses and placentas from in vitro-produced bovine embryos. Theriogenology, 65, 178–191. doi: 10.1016/j.theriogenology.2005.09.022

21. Farmer, J. L., Burghardt, R.C., Jousan, F. D., Hansen, P. J., Bazer, F. W., Spencer, T. E. (2008). Galectin 15 (LGALS15) functions in trophectoderm migration and attachment. FASEB Journal, 22, 548–560. doi:10.1096/fj.07-9308com

22. Freitag, N., Tirado-Gonzaĺez, I., Barrientos, G., Herse, F., Thijssen, V. L. J. L., Weedon-Fekjær, S. M., et al. (2013). Interfering with Gal-1-mediated angiogenesis contributes to the pathogenesis of preeclampsia. Proceedings of the National Academy of Sciences of the United States of America, 110, 11451–11456. doi: 10.1073/pnas.1303707110

23. Fukumori, T., Takenaka, Y., Yoshii, T., Kim, H. R. C., Hogan, V., Inohara, H., et al. (2003). CD29 and CD7 mediate galectin-3-induced type II T-cell apoptosis. Cancer Research, 63, 8302–8311. PMID: 14678989

24. He, J., Li, X., Luo, H., Li, T., Zhao, L., Qi, Q., et al. (2017). Galectin-3 mediates the pulmonary arterial hypertension–induced right ventricular remodeling through interacting with NADPH oxidase 4. Journal of the American Society of Hypertension, 11, 275–289. doi: 10.1016/j.jash.2017.03.008

25. Hughes, R. C. (1999). Secretion of the galectin family of mammalian carbohydrate-binding proteins. Biochimica et Biophysica Acta, 1473, 172–185. doi: 10.1016/s0304-4165(99)00177-4

26. Hyde, K. J., Schust, D. J. (2016). Immunologic challenges of human reproduction: an evolving story. Fertility and Sterility, 106, 499–510. doi: 10.1016/j.fertnstert.2016.07.1073

27. Lamb, G. C., Dahlen, C. R., Larson, J. E., Marquezini, G., Stevenson, J. S. (2010). Control of the estrous cycle to improve fertility for fixed-time artificial insemination in beef cattle: a review. Journal of Animal Science, 88 (13 Suppl), E181–E192. doi: 10.2527/jas.2009-2349

28. Liu, F. T., Patterson, R. J., Wang, J. L. (2002). Intracellular functions of galectins. Biochimica et Biophysica Acta, 572, 263–273. doi: 10.1016/s0304-4165(02)00313-6

29. Liu, F. T., Rabinovich, G. A. (2005). Galectins as modulators of tumor progression. Nature Reviews Cancer, 5, 29–41. doi: 10.1038/nrc1527

30. Machado, R., Corrêa, R. F., Barbosa, R. T., Bergamaschi, M. A. C. M. (2008). Escore da condição corporal e sua aplicação no manejo reprodutivo de ruminantes. EMBRAPA São Carlos, Brasil Circular Técnica 57. ISSN 1981-2086

31. Marques, M. O., Morotti, F., Lorenzetti, E., Bizarro-Silva, C., Seneda, M. M. (2018). Intensified use of TAI and sexed semen on commercial farms. Animal Reproduction, 15, 197–203. doi: 10.21451/1984-3143-AR2018-0070

32. Modenutti, C. P., Capurro, J. I. B., di Lella, S., Martí, M. A. (2019). The Structural Biology of Galectin-Ligand Recognition: Current Advances in Modeling Tools, Protein Engineering, and Inhibitor Design. Frontiers in Chemistry, 7, 823.

33. Moreira, R. (2019, january 01). Aumentar em 1% a taxa de prenhez reduz em 10% o custo da IATF. Giro do boi. https://www.girodoboi.com.br/videos/aumentar-em-1-na-taxa-de-prenhez-reduz-em-10-o-custo-da-iatf/

34. Nickel, W. (2005). Unconventional secretory routes: direct protein export across the plasma membrane of mammalian cells. Traffic, 6, 607–614. doi: 10.1111/j.1600-0854.2005.00302.x

35. Perillo, N. L., Pace, K. E., Seilhamer, J. J., Baum, L. G. (1995). Apoptosis of T cells mediated by galectin-1. Nature, 378, 736–739. doi: 10.1038/378736a0

36. Pohler, K., Peres, R., Green, J., Graff, H., Martins, T., Vasconcelos, J., Smith, M. (2016). Use of bovine pregnancy-associated glycoproteins to predict late embryonic mortality in postpartum Nelore beef cows. Theriogenology, 85, 1652–1659. doi: 10.1016/j.theriogenology.2016.01.026

37. Pope, W. (1988). Uterine asynchrony: a cause of embryonic loss. Biology of Reproduction, 39, 999–1003. doi: 10.1095/biolreprod39.5.999

38. Ramhorst, R. E., Giribaldi, L., Fraccaroli, L., Toscano, M. A., Stupirski, J. C., Romero, M.D., et al. (2012). Galectin-1 confers immune privilege to human trophoblast: implications in recurrent fetal loss. Glycobiology, 22, 1374–1386. doi: 10.1093/glycob/cws104

39. Reese, S. T., Franco, G. A., Poole, R. K., Hood, R., Fernadez-Montero, L., Oliveira Filho, R., et al. (2020). Pregnancy loss in beef cattle: A meta-analysis. Animal Reproduction Science, 212, 106251. doi: 10.1016/j.anireprosci.2019.106251

40. Rubisnstein, N., Ilarregui, J. M., Toscano, M. A., Rabinovich, G. A. (2004a). The role of galectins in the initiation, amplification and resolution of the inflammatory response. Tissue Antigens, 64(1):1-12. doi: 10.1111/j.0001-2815.2004.00278.x

41. Rubinstein, N., Alvarez, M., Zwirner, N. W., Toscano, M. A., Ilarregui, J. M., Bravo, A., et al. (2004b). Targeted inhibition of galectin-1 gene expression in tumor cells results in heightened T cell-mediated rejection: a potential mechanism of tumor-immune privilege. Cancer cell, 5, 241–251. doi:10.1016/S1535-6108(04)00024-8

42. Santos-de-Oliveira, R., Dias-Baruffi, M., Thomaz S. M., Beltramini, L. M., Roque-Barreira, M. C. (1994). A neutrophil migration-inducing lectin from Artocarpus integrifolia. The Journal of Immunology, 153, 1798–1807. https://www.jimmunol.org/content/153/4/1798

43. Sharon, N., Lis, H. (1986). Lectin biochemistry. New way of protein maturation. Nature, 323, 203–204. doi: 10.1038/323203a0

44. Sharon, N., Lis, H. (1989). Lectins as cell recognition molecules. Science, 246, 227–234. doi: 10.1126/science.2552581

45. Schäfer, T., Zentgraf, H., Zehe, C., Brügger, B., Bernhagen, J., Nickel, W. (2004). Unconventional secretion of fibroblast growth factor 2 is mediated by direct translocation across the plasma membrane of mammalian cells. Journal of Biological Chemistry, 279, 6244–6251. doi: 10.1074/jbc.M310500200

46. Stowell, S. R., Karmakar, S., Stowell, C. J., Dias-Baruffi, M., McEver, R. P., Cummings, R. D. (2007) Human galectin-1,-2, and -4 induce surface exposure of phosphatidylserine in activated human neutrophils but not in activated T cells. Blood, 109, 219–227. doi: 10.1182/blood-2006-03-007153.

47. Teichberg, V. I., Silman, I., Beitsch, D. D., Resheff, G. (1975). A beta-D-galactoside binding protein from electric organ tissue of Electrophorus electricus. Proceedings of the National Academy of Sciences, 72, 1383–1387. doi: 10.1073/pnas.72.4.1383

48. Than, N. G., Romero, R., Erez, O., Weckle, A., Tarca, A. L., Hotra, J., et al. (2008). Emergence of hormonal and redox regulation of galectin-1 in placental mammals: implication in maternal–fetal immune tolerance. Proceedings of the National Academy of Sciences, 105, 15819–15824. doi:10.1073/pnas.0807606105

49. Vasta, G. R. (2012). Galectins as pattern recognition receptors: structure, function, and evolution. Advances in Experimental Medicine and Biology, 946 s21–36. doi: 10.1007/978-1-4614-0106-3_2

50. Vergetaki, A., Jeschke, U., Vrekoussis, T., Taliouri, E., Sabatini, L., Papakonstanti, E. A., et al. (2014). Galectin-1 overexpression in endometriosis and its regulation by neuropeptides (CRH, UCN) indicating its important role in reproduction and inflammation. Plos One, 9, e114229. doi: 10.1371/journal.pone.0114229

51. Yang, R. Y., Hsu, D. K., Liu, F. T. (1996). Expression of galectin-3 modulates T-cell growth and apoptosis. Proceedings of the National Academy of Sciences, 93, 6737–6742. doi: 10.1073/pnas.93.13.6737

52. Wang, J. L., Gray, R. M., Haudek, K. C., Patterson, R. J. (2004). Nucleocytoplasmic lectins. Biochimica et Biophysica Acta, 1673, 75–93. doi: 10.1016/j.bbagen.2004.03.013

